# Delayed Trp53 activation protects *Dnmt3a*-mutant hematopoietic stem cells from inflammatory attrition

**DOI:** 10.1101/2025.01.27.635076

**Authors:** Ludovica Marando, George Giotopoulos, Pedro Madrigal, Dhoyazan M A Azazi, Ryan Asby, María Erendira Calleja-Cervantes, Muxin Gu, David Lara-Astiaso, Nicola K Wilson, Rebecca Hannah, Jaana Bagri, Emily Calderbank, Elisa Laurenti, Elsa Bernard, Sarah J Horton, Berthold Göttgens, Elli Papaemmanuil, Shuchi Agrawal-Singh, Malgorzata Gozdecka, George S Vassiliou, Brian J P Huntly

## Abstract

Hematopoietic stem cells (HSCs) accumulate somatic mutations over time, some conferring a fitness advantage that can lead to clonal hematopoiesis (CH). Mutations in *DNMT3A*, particularly at hotspot R882, are the most prevalent in CH and carry an increased risk of acute myeloid leukemia (AML). Although *DNMT3A* R882 mutations are linked to global DNA hypomethylation, the mechanisms underlying their selective advantage remain unclear. Here, we show that *Dnmt3a-R882H* mutant HSCs exhibit resilience under inflammatory and genotoxic stress. During IL-1β-induced emergency granulopoiesis, *Dnmt3a ^R882H/+^* HSCs uncouple increased proliferation from stem cell exhaustion. In contrast, wild-type HSCs rapidly progress to terminal differentiation. We link this phenotype to a delayed activation of the p53-p21-DREAM axis, that allows mutant HSCs to avoid attrition, despite increased replication stress. Similarly, mutant HSCs exhibit delayed Trp53 activation following irradiation, but eventually recover a physiological Trp53 response. Analysis of patient data reveals shared phenotypic features between *DNMT3A* and monoallelic *TP53* mutations in CH and myeloid neoplasms, highlighting potential functional similarities. Collectively, these findings suggest that the expansion of *DNMT3A*-mutant clones is affected by impaired TP53 signaling, which confers resilience against stressors. Therapeutic strategies targeting inflammatory pathways or the p53-p21-DREAM axis may reduce *DNMT3A-*CH expansion and/or progression and its associated risks.

## Introduction

Hematopoietic stem cells (HSCs) accumulate somatic mutations over time^1^. While most are inconsequential, some confer a fitness advantage that can eventually lead to a sizable clonal expansion, a phenomenon termed clonal hematopoiesis (CH)^2,3,4^. Mutations in *DNMT3A* account for more than half of all CH cases^2,3,4,5^. *DNMT3A* is also mutated across different Myeloid Neoplasms (MN), including Acute Myeloid Leukemia (AML)^6^, Myelodysplastic Syndrome (MDS)^7^ and Myeloproliferative Neoplasms (MPN)^8^. Mutations have been reported throughout the gene, but most commonly affect the arginine at codon-R882 in CH and at even higher relative frequencies in AML^9,10^, accordingly, *DNMT3A*^R882^ carriers have a ∼2-fold-increased risk of progression to AML^9^. Mutations at hotspot R882 produce a protein with dominant negative effects, resulting in global DNA hypomethylation^11^. *Dnmt3a* loss is associated with augmented HSC function^12,13,14^, however differences in methylation correlate poorly with differences in gene expression^15,16^. As a result, a unifying mechanism(s) that explains the fitness advantage of *DNMT3A*-mutant cells, as well as their propensity to AML progression, is currently lacking.

CH incidence increases with age^17^, but only progresses to MN in a fraction of cases^9^. An important risk factor for progression and CH-associated non-hematological disease is clonal size. As large clones carry a much greater risk than smaller ones^9,18,19,20^, inhibiting clonal expansion can have important benefits for CH carriers. However, efforts to limit clone size are hampered by our limited understanding of the mechanisms driving clonal expansion. The relative fitness advantage of *Dnmt3a-*mutant HSCs appears to be enhanced by inflammatory signaling via stressors such as TNF-α^21^, INF-γ^22,23^ and IL-6^24^, suggesting that preferential resilience to stress may be an important driver of *DNMT3A*-CH clonal expansion. However, the mechanistic basis for this remains unknown. Here, we characterize the behaviour of *Dnmt3a^R882H^*mutant HSCs/HSPCs under IL-1β-induced emergency granulopoiesis and radiation-induced stress. We report an altered function for *Dnmt3a^R882H^*mutant HSC at the molecular, cellular and tissue scale and link the competitive advantage to a blunted physiological Trp53 activation in response to these independent stressors. Using two large human datasets, we note that *DNMT3A^R882^*and *TP53* CH carry similar risks of evolving to MN over time and that MNs with either *DNMT3A*- or monoallelic *TP53*-mutations share multiple phenotypic similarities. These findings add to our understanding of the mechanisms driving *DNMT3A-CH* clonal fitness and identify potential strategies for mitigating its progression to a fully malignant state.

## Results

### *Dnmt3a^R882H/+^* cells show abnormal basal activation of inflammatory pathways

We and others have shown that *Dnmt3a*-mutant hematopoietic cells outcompete *Dnmt3a* wild-type cells under conditions of cellular stress^12,13,15,21,22,23,25^. Here, using a conditional knock-in mouse model (Fig.S1A), we interrogate these dynamic responses in cells heterozygous for *Dnmt3a^R882H^* (hereafter *Dnmt3a^R882H/+^*, mutant or mut) against their wild-type counterparts (hereafter *Dnmt3a^WT^* or WT).

Although *Dnmt3a^R882H/+^* and *Dnmt3a^WT^*bone marrow (BM) cells are similar immunophenotypically, the former show enhanced self-renewal *in vitro* (****p<0.001, Fig.S1B-C). Using scRNA-Seq (10x Genomics^26^), we characterized the transcriptome of single LK (lin-/c-kit+) BM cells at steady state (16,266 *Dnmt3a^WT^*, 14,776 *Dnmt3a^R882H/+^* cells, 2 mice/genotype) and assigned cellular identity to 13 clusters (Fig.1A). Mut and WT cells had a similar cellular composition and pseudobulk analysis revealed 391 genes differentially expressed between the two genotypes (adj.p.val.<0.05), with only 11 of these demonstrating an absolute fold-change (FC) of >1.5 (supplementary table 1). Notably, many of these were interferon (INF) response genes that were downregulated in mutant cells (Fig.S2B-C), a pattern also evident within the HSC-enriched cluster (cluster1) (Fig.1B-C, Fig.S2A, supplementary table 2).

**Figure 1.**
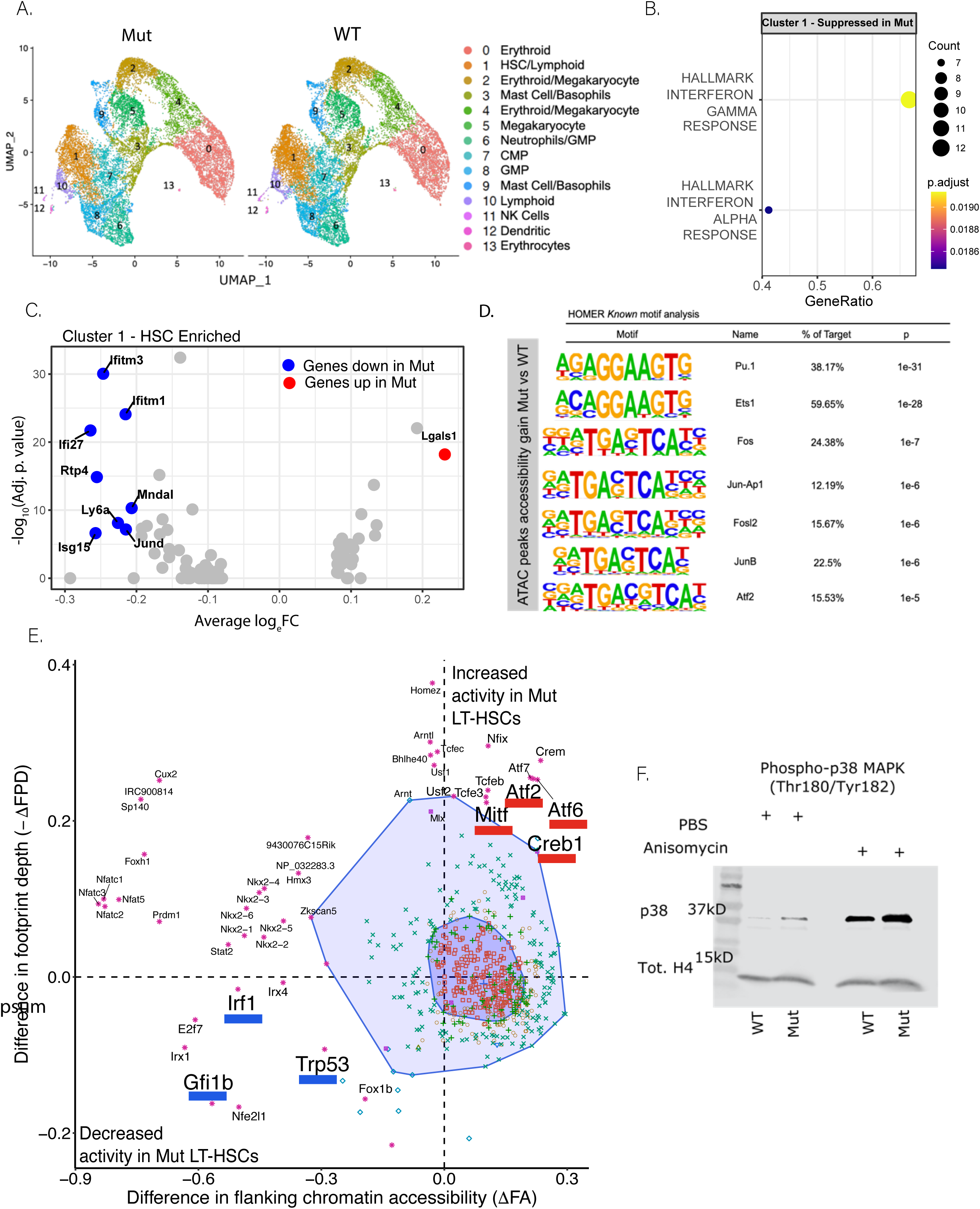
*Dnmt3a*^*R882H/+*^ cells show abnormal activation of inflammatory pathways at steady state. See together with Fig.S2 A. UMAP showing cluster distribution and cluster annotation of single LK BM cells from 2 mutant and 2 WT mice. B. Pseudobulk analysis of cluster1 - GSEA analysis of DEGs within the HSC-enriched cluster (cluster1) showing downregulation of interferon signalling genes in mutant cells. C. Volcano plot showing significant DEGs (adj.p.val.<=0.05 and absolute FC>1.25) within cluster1. D. Known motifs significantly enriched at genomic regions gaining accessibility in the presence of Dnmt3a^R882H/+^. E. BaGFoot analysis illustrates TFs with differential footprint depth and accessibility in Dnmt3a^R882H/+^ vs. WT LT-HSCs. Data points within bag-and-fence area (dark and light purple), including 50% and >97% of the population respectively, are not significant (NS). Motifs outside the fence and with a p value<0.05 (two-tailed, not multiple testing corrected) are statistically significant outliers. F. Western blot analysis of p38 MAPK phosphorylation at steady state, and upon induction of ribotoxic stress with anisomycin (positive control) in HSPC.

Given these modest transcriptional differences in homeostasis between the genotypes, but their altered function, we investigated whether DNMT3A loss-of-function^11,16,27^ could affect chromatin accessibility, facilitating differential responses to external stimuli. ATAC-Seq analysis of ESLAM LT-HSCs^28^, identified increased accessibility at 689 loci in mut LT-HSCs (including regions enriched for Pu.1, Ets and Ap-1-complex motifs, Fig.1D) and decreased accessibility at 1,556 loci (supplementary table 3). We then used Bivariate Genomic Footprinting (BaGFoot)^29^, to simultaneously assess differential footprint depth of transcription factors (TFs) and alterations in motif-flanking accessibility genome-wide. BagFoot analysis revealed decreased accessibility and reduced footprint depth of Interferon Regulatory Factor 1 (Irf1) in mutant LT-HSCs (Fig.1E). Of interest, Trp53, the mouse homologue of TP53, also demonstrated reduced binding in mutant cells, as did Gfi1b, a repressive TF regulating HSC quiescence^30^ (Fig.1E). Conversely, increased accessibility and footprint depth, indicating increased binding in mutant cells, was noted for TFs involved in the p38-MAPK signalling pathway (Atf2, Creb1, Mitf, Aft6 Fig.1E). Corroborating this finding, *Dnmt3a^R882H/+^* Lin^-^ cells showed higher basal p38 MAPK auto-phosphorylation at steady-state, a difference lost upon treatment with the potent activator of the p38/JNK-MAPK pathway, anisomycin (Fig.1F, Fig.S2D).

Taken together, our analyses suggest that differential basal activation of key inflammatory pathways in *Dnmt3a^R882H/+^* Hematopoietic Stem and Progenitor cells (HSPCs), may favour their clonal expansion under inflammatory stress and underlie CH-associated comorbidities^21,22,23,31,32,33,34^.

### *Dnmt3a^R882H/+^* LT-HSCs show an altered response to inflammatory signals

P38α MAPK is a pivotal cell-cycle and metabolic regulator of HSPCs upon hematological “stress”^35^, especially during emergency granulopoiesis^35^. To determine how basal signaling alterations in p38α MAPK influence stress responses, we exposed LT-HSCs to IL-1β, a key driver of emergency granulopoiesis^36^. Abnormal clonal expansion of *Dnmt3a^R882H/+^* HSCs has been reported following TNF-α-^21^, INF-γ-^22,23^ and IL-6-stimulation^24^, but the effects of IL-1β exposure were not previously investigated. In addition, increased IL-1β expression in monocytes of *DNMT3A* -CH patients with heart failure^37^ supports a potential paracrine role. We therefore assessed the division, proliferation and differentiation characteristics of *Dnmt3a^R882H/+^*LT-HSCs post-IL-1β-stimulation.

Single LT-HSCs from mutant and WT mice were cultured in media supplemented with IL-1β, TPO or IL-6 (Fig.2A). IL-1β- and IL-6-stimulation, led to a higher proportion of *Dnmt3a^R882H/+^*LT-HSCs completing first cell-division by 40h (**p=0.006 and *p=0.03 respectively Fig.2B), with the effects most marked for IL-1β, leading to us focusing on this stimulation. We then performed longitudinal analysis of the divisional kinetics of >1000 LT-HSCs across 7 biological replicates for each genotype, assessing the number of cell divisions at 10-12h intervals for the first 72h. Upon IL-1β-stimulation, *Dnmt3a^R882H^*^/+^ LT-HSCs exited quiescence earlier than WT cells (**p=0.0051 Fig.2C). A similar trend was observed between unstimulated conditions, suggesting that faster exit from quiescence might be an intrinsic *Dnmt3a^R882H^*^/+^ property accentuated by IL-1β-stimulation. Similarly, when the kinetics of exit from quiescence were studied *in vivo* (Fig.2D), we observed that the proportion of LT-HSCs in G1 and S-G2M plateaued at 12h in WT mice, whereas *Dnmt3a^R882H^*^/+^ LT-HSCs demonstrated a more prolonged exit from quiescence up to 24h (**p= 0.002 Fig.2E). Of note, in both genotypes, LT-HSCs returned to baseline quiescence levels by 36h post-IL-1β-stimulation.

**Figure 2.**
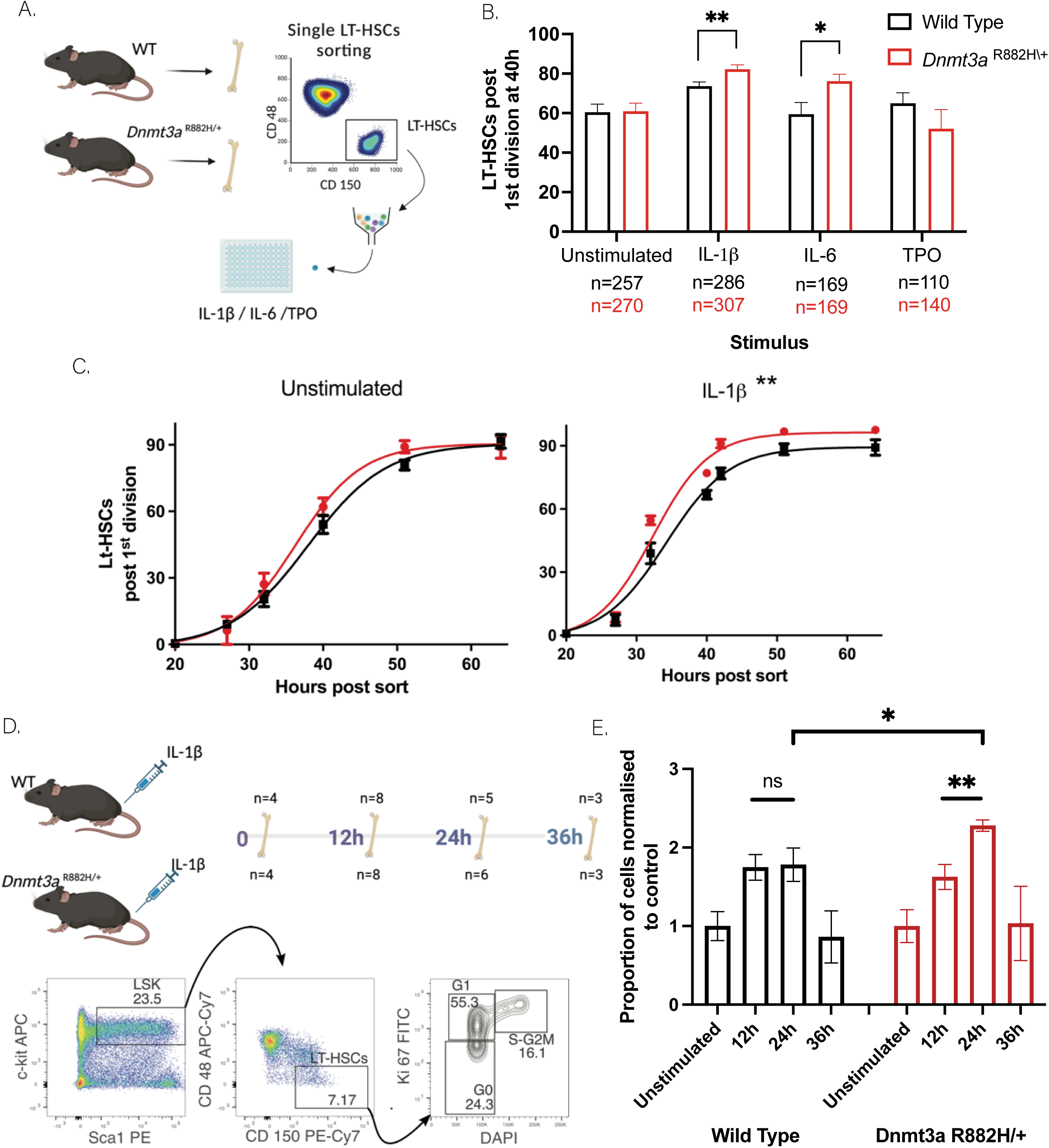
*Dnmt3a^R882H/+^*LT-HSCs show an altered response to inflammation. See together with Fig.S3 A. Single LT-HSCs from mutant and WT animals were sorted in 96-well plates and cultured with IL-1β-, TPO- or IL-6-enriched media. B. A higher proportion of Dnmt3a^R882H/+^ LT-HSCs completed first division by 40h upon IL-1β stimulation (82% vs. 75% **p=0.0067, unpaired t-test) and IL-6 (76% vs. 59% *p=0.033, unpaired t-test). n=number of single LT-HSCs observed in 7 mutant and 7 WT mice for PBS and IL-1β, 2 mutant and 2 WT mice for TPO and 4 mutant and 4 WT mice for IL-6, data is shown as mean±SEM). C. Sigmoid test fit modelling across the different time-points shows that Dnmt3a^R882H/+^ LT-HSCs exited quiescence early compared to WT following IL-1β stimulation (**p=0.0051, 2-way ANOVA). n=number of single LT-HSCs observed in 7 mutant and 7 WT mice across 5 independent experiments. D. Schema of the in vivo IL-1β stimulation experiment and gating strategy used to identify the different phases of the cell-cycle in LT-HSCs. E. Fold change proportion of LT-HSCs in cycle (G1 and S/G2-M), compared to the respective unstimulated control across the different time-points (**p= 0.002, *p = 0.039 unpaired t-test).

To further characterize the effects of IL-1β on LT-HSC behaviour, we assessed single-LT-HSC-derived clones by flow cytometry, following 7-10d in culture. Although clone viability and cell-death were similar between genotypes (Fig.S3A), mutant colonies contained significantly more live cells compared to WT colonies (**p=0.0061 Fig.3A). This suggests that the difference in the number of live events was secondary to a *bona fide* hyperproliferation, reflective of earlier entry into cell-cycle, of the *Dnmt3a^R882H^*^/+^ cells following exposure to IL-1β, rather than due to cell death of WT LT-HSCs.

**Figure 3.**
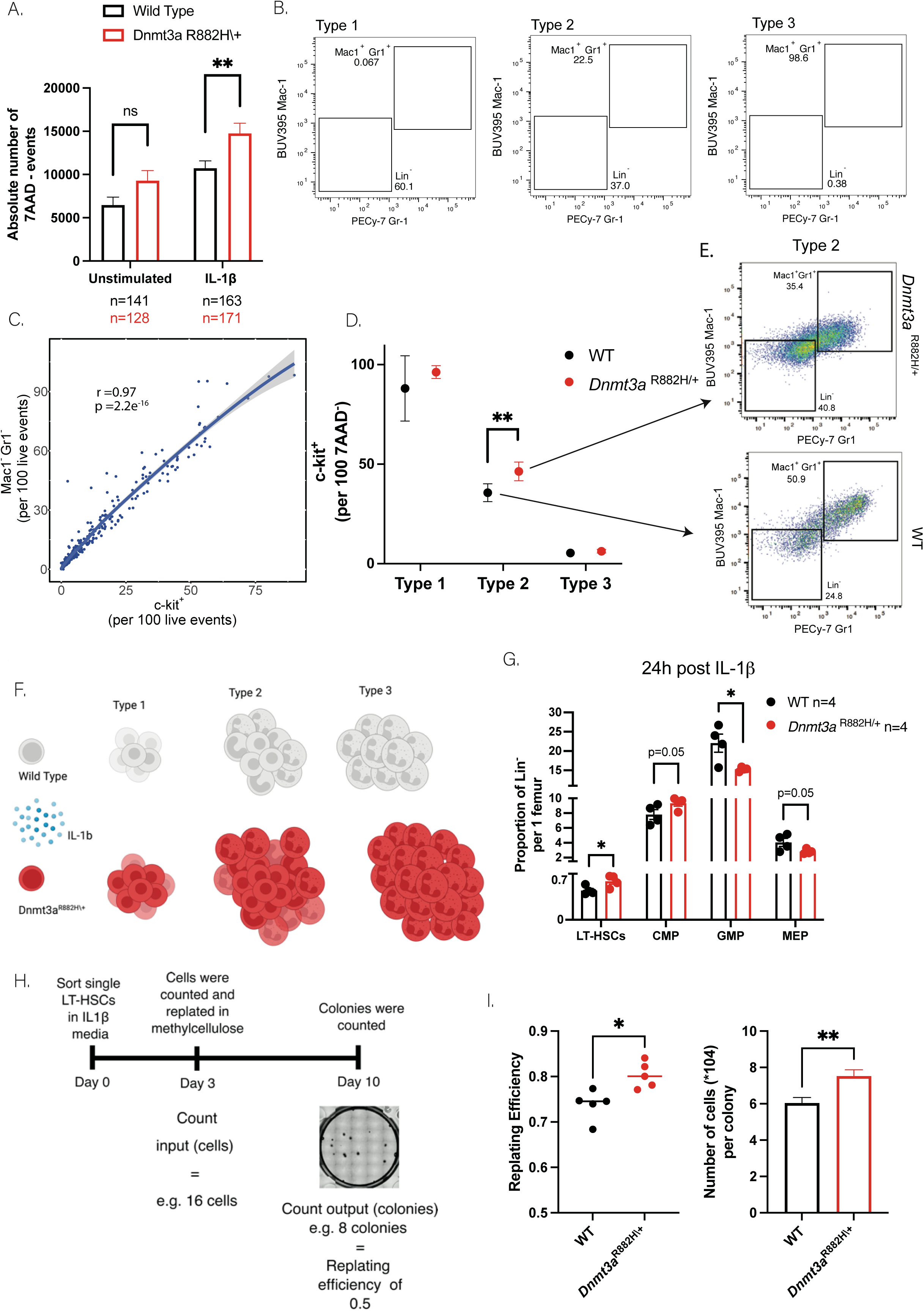
*Dnmt3a*^R882H/+^ LT-HSCs resist IL-1β-mediated depletion and are more proliferative under stimulation. See together with Fig.S4 A. Bar chart showing the absolute number of live events (7AAD^-^) enumerated with flow cytometry at 7-10d showing that IL-1β-treated Dnmt3a^R882H/+^ LT-HSCs give rise to colonies with a higher number of cells (**p=0.0061, unpaired t-test). Data are represented as mean±SEM. B. Representative examples of the observed single LT-HSC clonal output at 7-10d: Type 1 colonies (undifferentiated); Type 2 colonies (mixed); Type 3 colonies (fully differentiated). C. Linear correlation between the Mac1^-^Gr1^-^ gate and the c-Kit^+^ gate within the same colony (Pearson’s r=0.97). Ribbon represents the 5-95%CI. D. Dnmt3a^R882H/+^ type 2 colonies are qualitatively different from WT type 2 colonies and show a higher proportion of c-Kit^+^ cells (**p=0.0013, unpaired t-test). Data are represented as mean±SEM. E. Representative examples showing the qualitative differences observed in type 2 colonies. F. Schema of our model of emergency granulopoiesis in vitro. WT LT-HSC progress faster through terminal differentiation, whereas mutant LT-HSCs hyperproliferate whilst being able to maintain the size of their progenitor compartment for longer. G. Number of LT-HSCs and progenitors per femur were enumerated upon sacrifice at 24h post IL-1β administration, (*p=0.04 for LT-HSCs and *p=0.02 for GMP, unpaired t-test), data are represented as mean±SEM. H. Experimental layout and representative example of how replating efficiency was calculated. I. Dot plot showing replating efficiency across the 5 biological replicates per genotype (*p = 0.016, paired-t-test). Bar plot showing the average number of cells per colony (**p =0.001 shown, unpaired t-test). Data are represented as mean±SEM.

We then assessed differentiation outputs using lineage-specific markers. The clonal output of single LT-HSCs segregated into 3 different patterns (Fig.3B): Type-1 colonies (almost exclusively Mac1^-^Gr1^-^ cells, >80%), Type-2 (mixed output of Mac1^+^Gr1^+^&Mac1^-^Gr1^-^ cells), and, Type-3 (almost exclusively Mac1^+^Gr1^+^ cells, >80%). Type-1 colonies represented the most common state in unstimulated wells, whereas type-3 colonies represented the most common state in IL-1β-stimulated wells, regardless of genotype (Fig.S4A). Of note, Mac1^-^Gr1^-^ cells retained an immunophenotype that included both HSC (c-Kit^+^Sca-1^+^) and HSPC (c-Kit^+^Sca-1^-^) populations (Fig.3C, Fig.S4B), allowing us to capture proxies for different stages of myeloid maturation within the same colony. *Dnmt3a^R882H^*^/+^ LT-HSCs generated a higher number of both progenitors and differentiated cells (Fig.S4C). In addition, within *Dnmt3a^R882H^*^/+^ type-2 colonies we consistently identified a higher proportion of c-Kit^+^ progenitors compared to WT type-2 colonies (**p=0.0013, Fig.3D-F, Fig.S4D-E). To corroborate these findings *in vivo*, we enumerated LT-HSCs and c-Kit^+^ progenitors after 24h of IL-1β-stimulation, and observed greater retention of immature states, as well as larger numbers of LT-HSCs in *Dnmt3a^R882H^*^/+^ mice (*p=0.04 for LT-HSCs and *p=0.02 for GMP Fig.3G, Fig.S4F). Therefore, despite earlier cell cycle entry and higher proliferation rates, *Dnmt3a^R882H/+^* HSPCs retained a more primitive phenotype and appeared more resistant to IL-1β-mediated depletion. To assess the functional significance of these changes and link them to self-renewal, we assessed the clonogenic capacity of LT-HSCs following IL-1β-stimulation. Single LT-HSCs were plated and cultured for 3d with IL-1β and wells containing between 8-30 cells (ensuring at least three cell-divisions) were plated in methylcellulose (Fig.3H). Consistent with our immunophenotypic findings, *Dnmt3a^R882H/+^* cells retained better clonogenic efficiency and gave rise to colonies containing more cells (*p=0.016 and **p=0.001 respectively Fig.3I).

### IL-1β-stimulation induces altered gene expression and replicative stress in*Dnmt3a^R882H/+^* LT-HSCs

To gain mechanistic insight into the phenotypic differences between WT and *Dnmt3a^R882H/+^* LT-HSCs, we interrogated their transcriptomes at 12-, 24- and 36h-post IL-1β-stimulation (Fig.4A). At 12h, *Dnmt3a^R882H/+^* HSCs showed distinct gene expression profiles (Supplementary Tables 4-6) with 3124 differentially upregulated and 2765 downregulated genes [FC>1.5 and q>0.8^38,39^; Fig.4B]. Similar differences were observed at the other time-points, with 2080 and 3081 genes (at 24h) and 3256 and 2163 genes (at 36h) up- and down-regulated respectively (supplementary tables 4-6, Fig.S5A).

**Figure 4.**
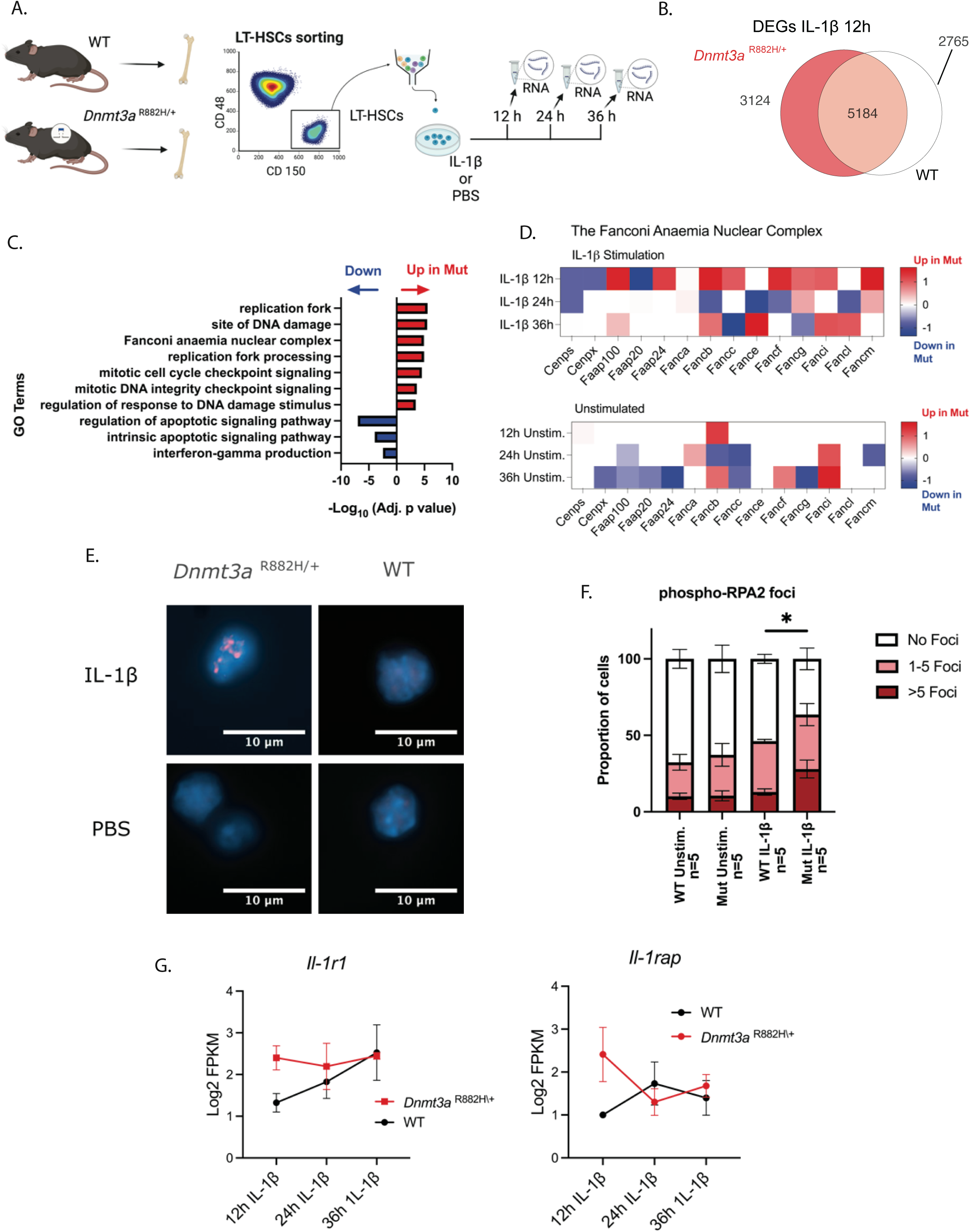
IL-1β-stimulation induces altered gene expression and replicative stress in *Dnmt3a^R882H/+^* LT-HSCs. A. Schema of the RNA-Seq experiment. B. Venn Diagram of DEGs at 12h (significant DEGs were defined as absolute FC>1.5 and q>0.8) C. Top GO terms for mutant IL-1β 12h vs. WT IL-1β 12h D. Heatmap showing expression of the components of the FA nuclear complex in mutant vs. WT cells in stimulated and unstimulated samples across all time-points. E. Representative images showing phospho-RPA2 nuclear localization via IF. F. Bar chart showing the proportion of cells with nuclear phospho-RPA2 foci in progenitors from WT and mutant mice, demonstrating an increase in the proportion of cells with >5 foci in Dnmt3a^R882H/+^ progenitors following stimulation with IL-1β (*p=0.033, unpaired t-test). Data are represented as mean±SEM. G. Gene expression as Log2 FPKM±SEM, for the IL-1β receptor complex across the time-points.

Gene set enrichment analysis (GSEA) revealed distinct expression patterns in *Dnmt3a*-mutant cells, especially at 12h post IL-1β-stimulation, with downregulation of apoptosis genes and upregulation of nuclear replication-fork stabilisation and DNA damage sensing genes (Fig.4C). In particular, increased expression of several Fanconi Anaemia (FA) nuclear complex genes was noted in mutant LT-HSCs at 12h (Fig.4D), suggesting increased IL-1β-mediated replication-fork stress, in line with the strong pro-proliferative IL-1β signals and our phenotypic observations. We therefore interrogated phospho-RPA2 nuclear localisation, a surrogate for replication fork stalling^40^, in Lin^-^ cells after 12h stimulation with IL-1β and observed significantly increased numbers of phospho-RPA2 foci in mutant Lin^-^ cells (*p=0.03 Fig.4E-F). Moreover, both IL-1-receptor1 (*Il1-r1*) and IL-1-receptor accessory protein (*Il1-rap*) were overexpressed in mutant samples at 12h (q=0.92 and q=0.94 respectively, Fig.4G), highlighting a possible mechanism that explains our findings.

### Transcriptional dysregulation of the TP53-DREAM pathway and enhanced G1/S transition in *Dnmt3a^R882H/+^*LT-HSCs upon IL-1β-stimulation

DNA damage-sensing by FANCM and FAAP24 activates an ATR kinase-dependent checkpoint response, leading to phosphorylation and activation of multiple FA proteins^41^ and TP53^42^ haltIing cell-cycle progression and allowing repair of stalled replication forks^43,44^ (Fig.5A). Of note and fitting with the single cell functional experiments, at 24h post-stimulation, *Dnmt3a^R882H/+^* LT-HSCs showed increased expression of genes associated with G1-S transition, including members of the HALLMARK E2F and MYC targets V1 gene sets (Fig.5B, Fig.S5B-C and supplementary table 5).

**Figure 5.**
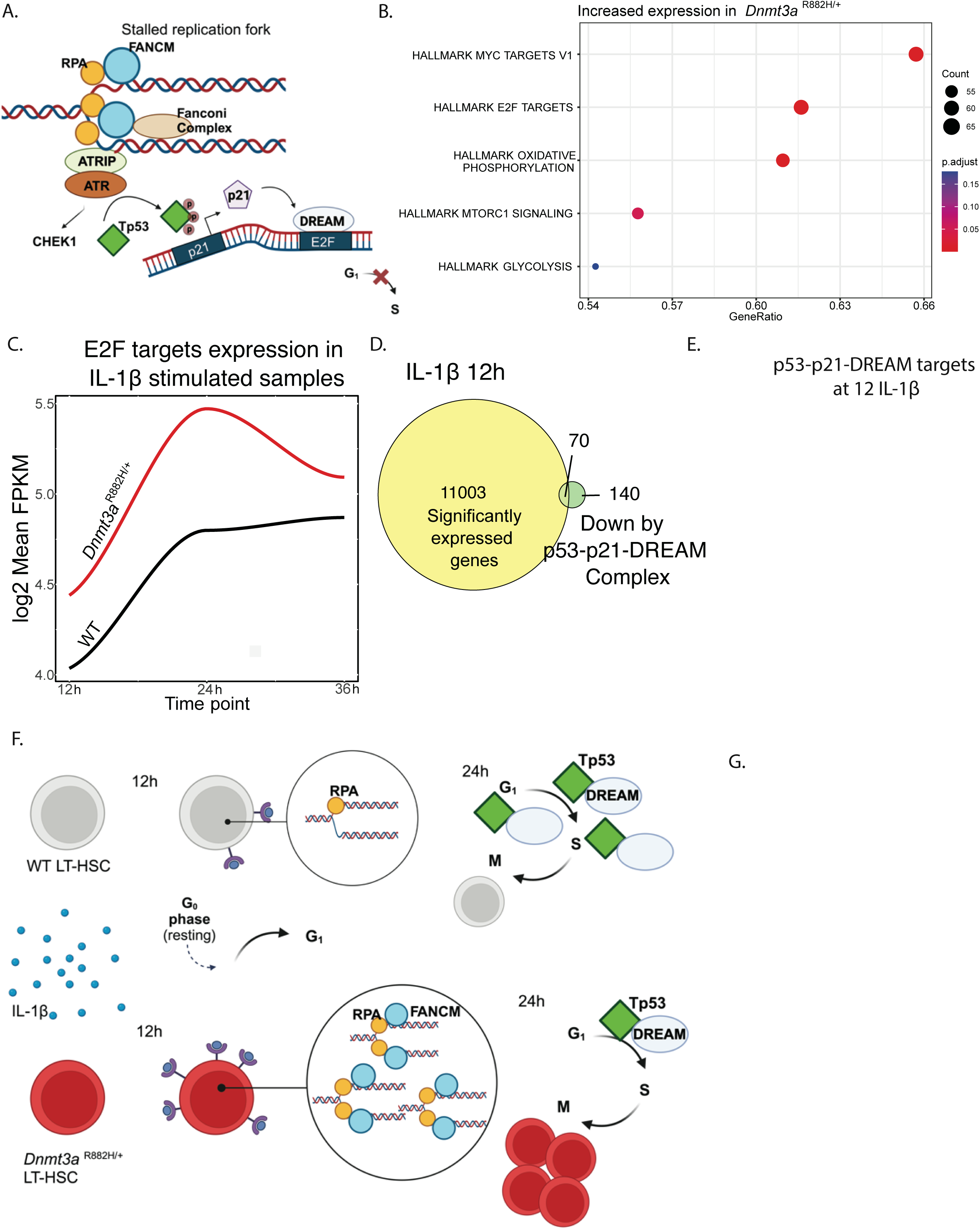
Transcriptomic analysis reveals dysregulation of the TP53-DREAM pathway and enhanced G1/S transition in *Dnmt3a^R882H/+^* LT-HSCs under IL-1β-stimulation. See together with Fig.S5&S6 A. Schema of the RPA/FA complex pathway leading to activation of ATR and TP53 mediated cell-cycle arrest, through the p21/DREAM axis B. GSEA analysis for significant DEGs (q>0.8) after 24h of IL-1β-stimulation. C. Kinetics of E2F activation, Log2 Mean FPKM expression of genes within the Hallmark E2F targets gene set (all genes showing signficant probability of differential expression, q>0.8, were included irrespective of FC) was compared between mutant and wild-type samples across the different time-points. LOESS regression analysis was used to fit a smoothed curve. D. Of the 210 p53-p21-DREAM-E2F/CHR targets identified by Fisher et al.^47^, we identified 70 genes in our 12h dataset (of 11,073 significant DEGs) that reached significant probability of differential expression (q>0.8, any FC). E. Gene expression as mean Log2 FPKM for the 70 p53-p21-DREAM-E2F/CHR targets previously identified, showing lack of repression of these genes in mutant LT-HSCs upon IL-1β-stimulation (p<0.0001 by paired t-test, mean and 5-95%CI are represented). F. Our proposed model of differential response to IL-1β. G. Bar chart showing that Niam siRNA induces hyperproliferation of sorted single WT LT-HSCs exposed to IL-1β, as shown by the increased number of live cells (7AAD^-^) (*p=0.0365, unpaired t-test). Data are represented as mean±SEM.

We next compared the normalized mean read count (FPKM) of all “HALLMARK E2F targets gene set” members between mutant and WT LT-HSCs (all genes with significant probability of differential expression, q>0.8, were included irrespective of FC) and used this gene set to infer the kinetics of E2F target activation over time. As expected, *Dnmt3a^R882H/+^* LT-HSCs exhibited a significantly earlier peak in E2F activity at 24 hours compared to wild-type cells (Fig. 5C). This observation aligns with the faster cell-cycle kinetics previously noted.

Given the reduced Trp53 TF activity in mutant cells (Fig.1E), we hypothesized that a delayed or abrogated Trp53 response could explain our findings. Upon DNA damage, TP53 facilitates cell-cycle arrest indirectly through *p21/CDKN1A,* and the transcriptional repressor complex DREAM^45,46^ (Fig.5A). Utilizing a 210 gene signature of p53-p21-DREAM targets^47^, we found that 70/210(∼33%) of these targets were differentially expressed (q>0.8, Fig.5D, supplementary table 7) within our 12h-IL-1β dataset. Supporting a dysregulation of the T53-DREAM pathway in mutant LT-HSCs, these genes lacked repression in *Dnmt3a^R882H/+^* LT-HSCs in comparison to WT cells (****p<0.0001, Fig.5E). Moreover, analysis of the unstimulated samples revealed that *Niam/Tbrg1*, a TP53 target and a component of the TP53–p21– DREAM–E2F/CHR pathway^48^, was consistently downregulated across all time-points in unstimulated mutant LT-HSCs (Fig.S6A-B). NIAM facilitates TP53-mediated activation of p21^48,49^, indirectly affecting cell-cycle progression (Fig.5F). Transient silencing of *Niam* in IL-1β-stimulated WT LT-HSCs phenocopied the hyperproliferative effect seen in *Dnmt3a^R882H^*^/+^ LT-HSCs (*p=0.03 Fig.5G), suggesting that impairment of the TP53–p21–DREAM pathway contributes to reduced cell-cycle arrest.

### Delayed Trp53 activation and altered DNA damage response in *Dnmt3a^R882H^***^/+^** cells following irradiation

We next interrogated how mutant and WT cells respond to radiation exposure, a stronger inducer of DNA-Damage and Trp53 activation (Fig.6A). Given the reduced basal Trp53 TF activity in *Dnmt3a^R882H^*^/+^ HSCs (Fig.1E), we assessed Trp53 genomic binding levels with CUT&RUN in Lin^-^ cells from 3 mutant and 3 wild-type animals at 2h post-irradiation. This was coupled with RNAseq at 6h post-irradiation (Fig.6A). 2834 regions were bound by Trp53 in WT samples upon irradiation (supplementary table 8). Analysis of signal enrichment across these regions, revealed reduced Trp53 binding-signal in *Dnmt3a^R882H^*^/+^ HSPCs under unirradiated and irradiated conditions (Fig.6B). This enrichment was particularly noted at genes involved in the Trp53 pathway/network (contained within the HALLMARK TP53 pathway gene set, Fig.6C), including exemplar genes *Cdkn1a, Bax, Fas* and *Mdm2*(Fig.6D). No difference was noted across samples for global Trp53 signal enrichment (Fig.S7A), supporting a selective reduction in Trp53 binding at its targets.

**Figure 6.**
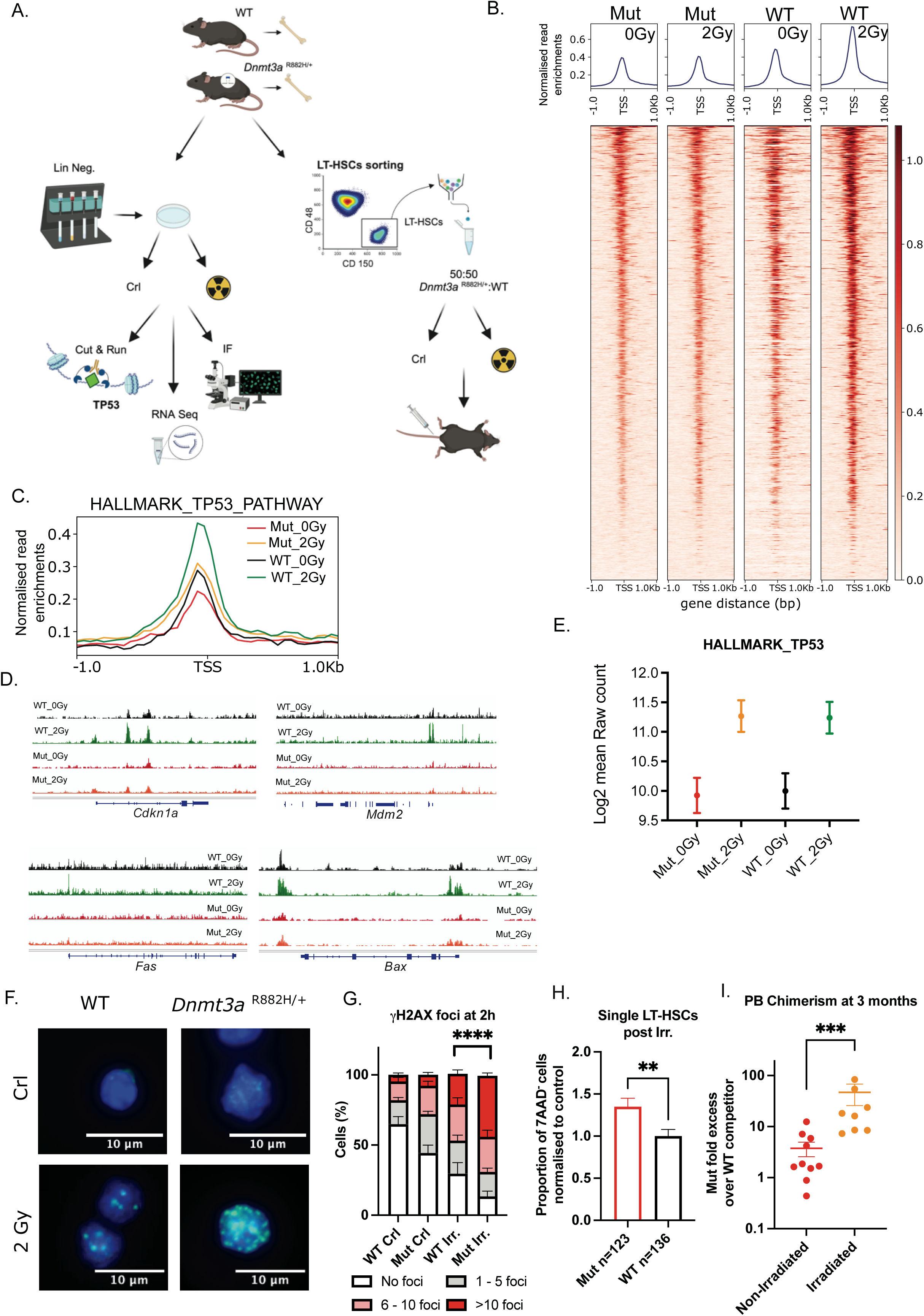
Delayed Trp53 activation and altered DNA damage response in *Dnmt3a^R882H^*^/+^ hematopoietic cells. See together with Fig.S7-S8 A. Experimental schema. Lin^-^ progenitors were irradiated (or sham-treated) ex vivo and subsequently underwent either Cut&Run (n=3 per genotype, 2h post-irradiation) or RNA-Seq (n=3 per genotype, 6h post-irradiation) or immunofluorescence (IF) for γH2AX foci enumeration (n=8 per genotype, 2h post-irradiation). LT-HSCs were irradiated (or sham treated) and immediately transplanted into lethally irradiated recipients (n=9 for each arm). B. Heatmap showing genomic enrichment scores for Trp53 binding, determined by CUT&RUN, in a ±1kb window from the transcription start site (TSS) of 2834 genes bound to Trp53 in WT samples upon irradiation. Top plot shows total signal enrichment at each bin for all replicates belonging to their respective condition (Mutant: unirradiated and irradiated, plus WT: unirradiated and irradiated). Each row of the heatmap corresponds to a gene in the 2834 gene set, sorted ascendingly by the signals from the mutant unirradiated samples. Genes where no signal was recorded for this set of samples were deemed as zeros, and excluded from the heatmap. C. Trp53 signal across the genes contained within the HALLMARK TP53 PATHWAY gene set, displayed are peaks 1 kb up- and down-stream of TSS. D. Genome browser tracks showing CUT&RUN signal at Cdkn1a, Mdm2, Fas, Bax. E. Log2 mean raw counts for significant DEG at 6h post irradiaiton (adj.p.val.<=0.05) contained within the HALLMARK_TP53 Targets gene set. F. Representative immunofluorescence image of γH2AX foci in unirradiated and irradiated cells. G. Bar chart showing γH2AX foci, in progenitors from WT and Mut mice, 2h post irradiation. The mean±SEM is shown, n=8 independent experiments (**** p< 0.001 by unpaired t test). H. Bar chart showing proportion of 7AAD^-^ cells relative to unirradiated control, in colonies of irradiated mutant and wild-type LT-HSCs cultured in the presence of IL-1β for 7-10d (**p= 0.006 by unpaired t test). The mean±SEM is shown, n=8 independent experiments. I. Dot plot showing fold excess of mutant cells over WT competitors in the PB of irradiated (n=9) and non-irradiated (n=9) in vivo recipients (***p = 0.0002, Mann-Whitney test). The mean±SEM is shown.

RNA Seq analysis revealed upregulation of 1216 genes in WT and 1166 genes in mutant cells (adj.p.val.<0.05 Fig.S8A and supplementary tables 9-10) at 6h post-irradiation. Despite the reduced Trp53 binding in mutant progenitors at 2h post-irradiation, Trp53 target genes reached similar expression to wild type at 6h (Fig.6E), suggesting a delay, rather than complete loss, of Trp53 activation in mutant cells. Consistent with a delay in Trp53 activation, mutant HSPCs exhibited significantly higher levels of DNA damage shortly after irradiation, as indicated by increased γH2AX foci (a surrogate for double-strand breaks) at 2 hours post-irradiation (****p < 0.001, Fig. 6F-G) albeit that they retained a comparable clonogenic output to wild-type cells (Fig. S8B-C). However, by 24 hours, γH2AX foci were signficianctly reduced (*p=0.02, Fig. S8D), indicating that mutant HSPCs can resolve DNA damage over time. To assess functional outcomes at a later time-point, we sorted LT-HSCs from mutant and WT animals and observed lower levels of apoptosis in mutant LT-HSCs at 24h post irradiation (p=0.03* Fig. S8E), and when single LT-HSCs were plated in the presence of IL-1β following *ex-vivo* exposure to irradiation, mutant LT-HSCs retained their proliferative advantage (**p=0.006, Fig.6H). Similarly, irradiation substantially enhanced the advantage of *Dnmt3a^R882H^*^/+^ LT-HSCs in *in vivo* competitive transplantation assays, resulting in significantly higher mutant chimerism in recipient animals (***p<0.001, Fig6I).

### *TP53-*monoallelic and *DNMT3A^R882^*mutations share similar MN evolution rates, genetic landscape and clinical outcomes

Despite being relatively common individually, *DNMT3A*and *TP53* mutations rarely co-occur in MN^6,50^. Biallelic TP53 loss-of-function mutations strongly associate with copy number alterations (CNA) and structural chromosomal aberrations^51^, features not typically seen in *DNMT3A*-mutant MN^10^. This apparent contrast led us to investigate potential differences and similarities between these recurrent drivers. However, there is limited comparative data available to assess these mutations at the CH stage. To address this gap, we used large-scale population-level data from 454,340 UK Biobank participants, where CH individuals were identified from whole exome sequencing (WES), as previously described^9^. We assessed the odds ratio (OR) of developing AML or MDS for carriers of common CH mutations, and found that individuals with *DNMT3A^R882^* and *TP53* CH had a similar risk of developing AML or MDS compared to controls (*DNMT3A^R882^* OR=12.38 95%CI (8.96-16.72), TP53 OR=8.67 95%CI (3.71-17.31), Fig. 7A and Supplementary table 11). Given the pacuity of comparative CH data, we extended our analysis along the MN continuum, focusing on MDS and AML to further explore the clinical and biological relationships between these recurrent drivers. We analyzed clinical and genetic data of 3,323 treatment-naïve MDS individuals (Table 1)^52,53^ and identified 249 patients with *T P 5 3*multi-hit, 120 with *T P 5*-m*3*onoallelic and 106 *DNMT3A^R882^* patients with wild-type *TP53* (Fig.7B and supplementary tables 12-13). The median overall survival (OS) was 10.8 months (95%CI, 9.4-12.3) for *TP53* multi-hit, 24.6 months (95%CI, 18.6-30.7) for *DNMT3A^R882^* and 31.4 months (95%CI, 23.9-39) for *TP53* monoallelic cases (Fig.7C). Time-to-leukemia development was 9.9 months (95%CI, 8.4-11.4) for *TP53* multi-hit, 21.7 months (95%CI, 16.1-27.3) for *DNMT3A ^R882^* and 31.8 months (95%CI, 23.7-40) for *T P 5* m*3* onoallelic cases (Fig.7D). The median number of chromosomal aberrations, including CNA (but excluding chromosome 17 abnormalities that can also define *TP53* status), was 1 in *TP53* monoallelic (IQR=0-2), 0 in *DNMT3A^R882^* (IQR=0-1) and 8 in *TP53* multi-hit cases (IQR=5-10; Fig.7E), in agreement with the fact that even a single TP53 intact copy is enough to generally preserve genome integrity^52^. We also observed that *TP53* multi-hit cases required fewer cooperating mutations; the median number of additional single oncogenic gene mutations was 1 in *TP5 3*multi-hit (IQR=0-2), 2 in *TP53* monoallelic (IQR=1-4) and 3 in *DNMT3A^R882^* cases (IQR=2-5; Fig.7F). The pattern of co-mutations was also very similar between *TP53* monoallelic and *DNMT3A^R882^* cases (Fig.7G, Fig.S9A-B), but both were distinct from *TP53* multi-hit. Overall, we demonstrate that MN patients with a *TP53* monoallelic mutation show analogous co-mutation patterns and comparable clinical outcomes to *DNMT3A^R882^* MN patients, as opposed to *TP53* multi-hit patients that represent an entirely separate entity, associated with pronounced genomic instability, a lack of significant point mutations and dismal OS. These results demonstrate phenotypic similarities between *DNMT3A^R882^* and monoallelic *TP53* alterations and align with our broader hypothesis that subtle disruptions in *TP53* signaling, as observed in *Dnmt3a^R882H^* HSCs, can enable their persistence under inflammatory stress to eventually achieve clonal dominance.

**Figure 7.**
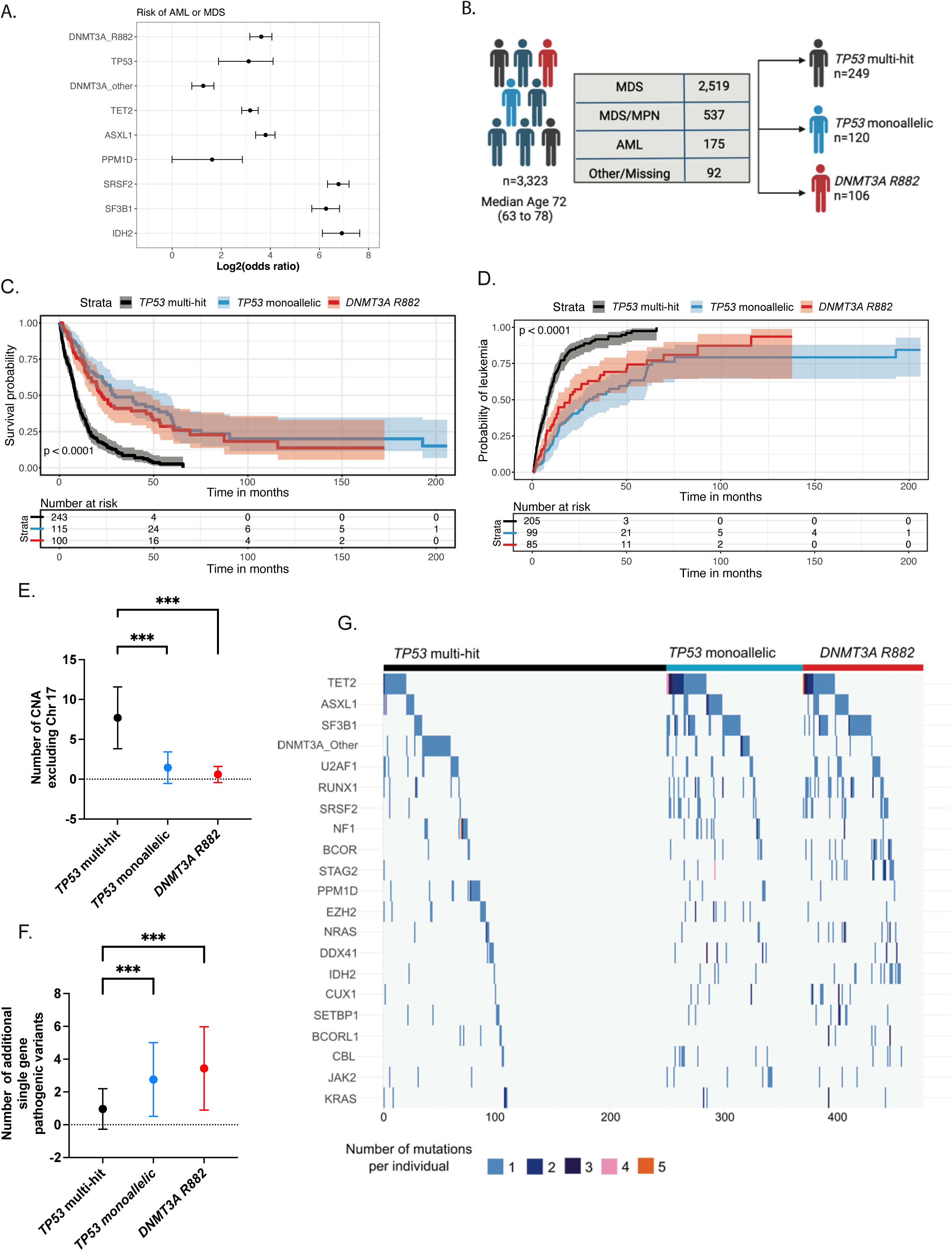
*TP53* monoallelic and *DNMT3A R882* myeloid neoplasms share similarities in their genetic landscape and clinical outcomes. See together with Fig.S9 A.Forest plot of the OR of AML or MDS per recurrent driver of CH within 454,340 UK Biobank participants, all drivers (except PPM1D) give a significant risk of AML or MDS p<0.0001 by fisher’s exact test, whiskers represent the 95% CI. B. Schema of the patient cohort. C. Kaplan-Meier probability of overall survival and D. cumulative AML incidence per group. p values were derived from two-sided log-rank test. E. Number of unique chromosomal alterations other than on Ch 17 per group. In addition to karyotyping, chromosomal aberrations were also detected on NGS sequencing and annotated by Bernard et al. ^26,27^. (****p<0.0001, Wilcoxon signed-rank test). Data is shown as mean± SD. F. Number of additional oncogenic driver per group. (****p<0.0001, Wilcoxon signed-rank test). Data is shown as mean±SD. G. Waterfall plot of mutation profiles in 249 patients with TP53 multi-hit, 120 patients with TP53 monoallelic and 106 patients with DNMT3A R882 MN. Each column represents a different individual. Only the top 21 most recurrent mutations are represented

**Table 1.** Characteristics of the study cohort. 1Q: first quartile; Q3 third quartile.

| Characteristics | Number of cases (%) | Median (1Q-3Q) |
| --- | --- | --- |
| **Gender** |  |  |
| Male | 2004 (60%) |  |
| Female | 1319 (40%) |  |
| **Age at diagnosis** |  | 72 (63 – 78) |
| Missing data | 20 (0.6%) |  |
| **WHO 2016 classification** |  |  |
| **MDS** |  |  |
| MDS Del 5q | 142 (4.3%) |  |
| MDS-EB1 | 464 (14%) |  |
| MDS-EB2 | 438 (13%) |  |
| MDS-SLD/MLD | 922 (28%) |  |
| MDS-RS-SLD/MLD | 467 (14%) |  |
| MDS-U | 86 (2.6%) |  |
| **AML** | 175 (5.3%) |  |
| **MDS/MPN** |  |  |
| CMML | 399 (12%) |  |
| aCML | 38 (1.1%) |  |
| MDS/MPN-RS-T | 100 (3%) |  |
| **Other** | 5 (0.15%) |  |
| **Missing data** | 87 (2.6%) |  |

## Discussion

CH-mutations arise across a range of ages, expand at different rates, and are influenced by differing selective processes and pressures^1,54,55^. A unifying and pervasive driver appears to be chronic inflammation, and previous studies have interrogated pro-inflammatory signalling (TNFα and INF-γ) in the context of *Dnmt3a* loss/mutation. Here, we expand on this by investigating the effects of IL-1β-stimulation on HSC quiescence, proliferation and differentiation. *Dnmt3a^R882H/+^* HSCs are more resistant to IL-1β-mediated depletion both *in vivo* and *in vitro*, demonstrating an earlier exit from quiescence, increased proliferation and a larger output of both HSPC and terminally differentiated cells. Mechanistically, we show that *Dnmt3a^R882H/+^* HSCs bypass replication stress-induced G1 arrest through a delay in the activation of the p53-p21-DREAM pathway, which permits hyperproliferation and clonal dominance in both short-term and long-term functional experiments.

Similarly, following irradiation, we demonstrate delayed Trp53 binding to target genes. This transient delay in Trp53 activation is sufficient to confer a clonal advantage while avoiding the long-term genomic instability typically associated with *TP53* multi-hit states. We demonstrate that *Dnmt3a^R882H/+^*HSCs eventually recover physiological levels of Trp53 activation, distinguishing their phenotype from the persistent dysfunction observed in biallelic TP53 alterations.

Our analysis of the UK Biobank cohort identifies that carriers of *DNMT3A^R882^* and *TP53* CH have a comparable risk of developing AML and MDS. In addition, CH population studies have also demonstrated closely matched estimates of gene fitness and clonal behaviour between *DNMT3A^R882^* and *TP53*-mutant HSCs^54,56^ providing insights into the role of these drivers in early disease evolution. Further corroboration of the link between *DNMT3A^R882^* and subtle *TP53* dysfunction also occurs further along the MN continuum, where analysis of a large human MDS cohort demonstrated that *DNMT3A* R882 and monoallelic *TP53* MDS share similarities in co-mutation patters and clinical outcomes. While these findings do not directly establish a shared functional mechanism, taken together with our experimental results, they suggest that *DNMT3A R882* mutations may partially mimic aspects of monoallelic *TP53* dysfunction, ultimately converging on similar trajectories of disease evolution. Additional evidence linking *DNMT3A*-mutation to altered TP53 function is also reported in a recent study showing that *DNMT3A^R882H^* preleukemic HSCs accumulate TP53 in a pseudo-mutant conformation^57^. Furthermore, similarly to *DNMT3A* CH and other CH genotypes, genomic instability is not the only mechanism of *TP53*-mutation mediated clonal advantage, and factors such as LPS-mediated release of IL-1β in murine models and chronic inflammation in patients^58^ can affect the clonal behaviour of TP53-mutant hematopoietic cells.

Our findings are in line with reported HSC phenotypes in *Trp53*-null mice^59^, showing increased HSC cycling, enhanced BM reconstitution capacity, elevated γH2AX foci and decreased apoptosis upon irradiation, albeit that *Trp53*-null rather than depleted cells demonstrated a more marked anti-apototic phenotype compared to wild-type, highlighting the phenotypic effects of precise Trp53 quantitation. In contrast to a recent study reporting increased Trp53 stabilisation and enhanced quiescence in *Dnmt3a^KO^*HSCs under INF-γ-stimulation^22^, our studies were conducted in a *Dnmt3a^R882H/+^*-mutant, rather than a *Dnmt3a*-null background, a genotype rarely seen in human CH. Possible critical differences between *DNMT3A R882*-mutations and complete loss of *DNMT3A* have also been reported by others^60,61^ and could account for these differing phenotypes.

In summary, our work associates the clonal advantage of *Dnmt3a^R882H/+^*HSCs, to an altered responsiveness to stressors such as inflammation and irradiation. These effects, particularly on Trp53 signalling, affect the cell cycle properties of mutant HSCs and are likely to account for selective advantages in CH, and have potential clinical implications. Several clinical trials are targeting inflammatory pathways in MN (using Canakinumab <u>NCT04239157</u>, <u>NCT05641831</u> and NLRP3 inhibitors <u>NCT05552469</u>, <u>NCT05552469</u>). Moreover, a preclinical study reported that anti-IL-6 antibodies reduce the selective advantage of *DNMT3A^mut^*HSCs under age-related fatty BM conditions^24^. Our study adds Trp53 and the DREAM complex to the list of potential therapeutic targets to modulate progression towards MN. A number of strategies, many in current clinical trials, exist to augment WT TP53 activity via targeting the MDM2-TP53 axis^46^. Modulation of the DREAM complex is also possible (e.g. through the use of CDK4/6 inhibitors such as Palbociclib^46^), should augmenting the other functions of TP53 give rise to unwanted side effects.

## Methods

### Mice

The Mx1-cre;*Dnmt3a^R882H^* mouse has been previously described (Fig.S1A)^25^. Unless otherwise specified, ∼6-month-old animals after a minimum of ∼4 months post Mx1-Cre-activation were used. Experiments were conducted under a UK Home Office project license following ethical review by the University of Cambridge Animal Welfare and Ethical Review Body.

### Isolation of cells, flow cytometry and cell sorting

Lineage negative (Lin^-^) cells were isolated using the EasySep^TM^ Mouse Hematopoietic Progenitor Cell Isolation Kit and subsequently stained using antibody cocktails (supplementary tables 14-18). Cell sorting/analysis was performed on a High-Speed Influx cell Sorter and BD LSRFortessas using FACSDiva and FlowJo.v9/v10.

### Long Term-Hematopoietic Stem Cell culture

Single CD45^+^EPCR^+^CD48^-^CD150^+^(ESLAM) Long Term-Hematopoietic Stem Cells (LT-HSCs) were sorted in 96-well plates and cultured in StemSpan^TM^ containing Pen/Strep, 2mM L-Glutamine, 0.1mM beta-Mercaptoethanol, 150ng/mL mSCF, 20ng/mL IL-11, 10%FBS ±25ng/mL IL-1β. At 7-10d, wells were stained with the appropriate antibody cocktail (supplementary table 16).

### Single-cell RNA Seq

Lineage c-Kit^+^ (LK) cells (supplementary table 18) from two wild-type and two mutant mice were processed using the Chromium^TM^ Single Cell 3’Library & Gel Bead Kit v2(10x Genomics).

### Bulk RNA of IL-1β stimulated LT-HSCs

∼500 LT-HSCs from two mutant and two wild-type mice were sorted on a 96-well plate (supplementary table 18). ‘Stimulated’ wells were treated with 25ng/mL IL-1β. RNA was extracted at 12h, 24h and 36h of culture, using the Arcturus^TM^ PicoPure^TM^ RNA Isolation Kit.

### RPA and H2AX immunofluorescence

Lin^-^ cells were cultured for 12h (±25ng/mL IL-1β), fixed and stained with anti-RPA2 pSer33(1:500) and AlexaFluor594 goat anti-rabbit(1:500).

For H2AX staining, Lin^-^ cells were cultured overnight in RPMI(20%FBS+IL-3+IL-6), irradiated (2Gy of X-rays) and returned to culture for 2h. Cells were stained using Histone H2AX-Ser139 (1:500) and AlexaFluor488 anti-mouse(1:500). A minimum of 100 cells per condition were scored using a Zeiss AxioSkop 2 microscope. Images were acquired using a Zeiss AxioImager Z2 and processed in ZeissZEN.

### Bulk RNA and CUT&RUN of irradiated samples

Lin^-^ cells from 3 wild-type and 3 mutant mice were cultured overnight and irradiated (2Gy of X-Rays). At ∼2h and ∼6h post-irradiation respectively, ∼500,000 cells from each sample were used for CUT&RUN (total-p53 antibody,#ab246223) or harvested for RNAseq.

### Competitive transplantation following irradiation of LT-HSCs

ESLAM LT-HSCs from three mutant and three wild-type animals received 1.25Gy of X-rays and a 50:50 mutant/wild-type mix of ∼400 LT-HSCs were injected into lethally irradiated congenic recipients.

### UK Biobank Analysis

UKB is a large-scale biomedical database and research resource containing genetic, lifestyle and health information from half a million UK participants.

Whole-exome sequencing of blood DNA from 454,340 UKB participants was used to identify somatic mutations using Mutect2 software as previously described^9^.

### Analysis on human samples

The MDS IWG cohort^52,53^ (https://www.cbioportal.org) was used. Eight cases with *TP53/DNMT3A ^R882^*co-mutation were excluded. Analysis was performed on 249 individuals with *TP53* multi-hit, 120 individuals with *TP53* monoallelic and 106 cases with *DNMT3A^R882^* and *TP53* wild-type. Additional data are provided in supplementary tables 12-13.

### Data sharing statement

All relevant data have been deposited in the Gene Expression Omnibus under GSE227026 (10x data), GSE275416 (LT-HSC ATAC-seq data), GSE275415 (LT-HSC RNA-seq data), GSE275414 (IR RNA-seq data) and GSE275218 (CUT&RUN data). This study does not report any original code.

## Acknowledgments

Funding in the laboratory of B.J.P.H. came from Cancer Research UK (CRUK) (C18680/A25508 and DRCRPG-Nov22/100014), the European Research Council (647685), the Medical Research Council (MRC) (MR-R009708-1 and MR-X008371), the Kay Kendall Leukaemia Fund (KKL1243), the ”La Caixa” Foundation (LCF/PR/HR20/52400016), The Leukemia & Lymphoma Society (7035-24), the CRUK Cambridge Centre (C49940/A25117 and CTRQQR-2021\100012) and the NIHR Cambridge Biomedical Research Centre (BRC-1215-20014) and was funded in part by the Wellcome Trust, which supported the Cambridge Stem Cell Institute (203151/Z/16/Z) and Cambridge Institute for Medical Research (100140/Z/12/Z). L.M. was funded by the Wellcome Trust PhD Programme for Clinicians (205254/Z/16/Z). J.B. is funded by the Wellcome Trust’s 4-Year MRes + PhD Programme in Stem Cell Biology and Medicine (218481/Z/19/Z). G.S.V. was a CRUK Senior Cancer Research Fellow (C22324/A23015).

The funders had no role in study design, data collection and analysis, decision to publish or preparation of the manuscript. The views expressed are those of the authors and not necessarily those of the NIHR or the Department of Health and Social Care.

## Author contributions

L.M, G.G & B.J.P.H. conceived the study, designed the experiments and wrote the manuscript. LM & GG designed and performed experiments, collected and interpreted data. P.M., D.M.A.A M.E.C-C and MG performed bioinformatic analysis and bioinformatic data visualisation. R.A. provided technical expertise and performed experiments. D.L-A performed ATAC-seq experiments. N.K.W. performed 10x experiments. R.H. processed 10x data. J.B. performed experiments. EC and EL provided technical help with single cell functional LT-HSC assays. E.B. & E.P. provided human patient sample data. S.A-S performed CUT&RUN experiments and interpreted data. M.G., B.J.P.H. & G.S.V. generated the *Dnmt3a^R882H^* mouse model. P.M., N.K.W., S.J.H., B.G., M.G. & G.S.V. reviewed and commented on the manuscript. All authors read the manuscript. B.J.P.H. supervised and directed the project.

## Materials & Correspondence

B.J.H.P.

## Supplemental information

Supplemental Document containing supplementary methods, figures S1–S9 and supplemental tables 13–18.

